# Social and ecological factors associated with innovation in urban sulphur-crested cockatoos (*Cacatua galerita*)

**DOI:** 10.64898/2025.11.30.691464

**Authors:** Lisa Fontana, Sofia Bolcato, Julia Penndorf, Lucy M Aplin

## Abstract

Why some species thrive in urban environments while others do not is a central question in behavioral ecology. Behavioral innovations has been proposed as a key mechanism facilitating this adaptation. At the individual level, innovativeness varies with cognitive and behavioral traits. However, at the population level, innovation rates can also be influenced by social and ecological factors including group size, and environmental novelty and complexity. The role of these factors are still under-explored, especially at within-city scales. To disentangle factors influencing group-level variation in innovation rates, we presented roosts of wild sulphur-crested cockatoos *Cacatua galerita*, with extractive-foraging tasks that required innovative problem-solving. We installed three tasks of different levels of difficulty on trees at fifteen communal roost sites across an urban matrix. We matched these with direct measures of roost-size and connectivity, and with high-resolution remote-sensing mapping to estimate variation in urbanization and environmental heterogeneity. We found that approach time was significantly associated with urbanization, with individuals in more urban sites approaching tasks more quickly, suggesting reduced neophobia with urbanization or increased familiarity with human-derived objects. In contrast, time to innovate in our study was explained by task difficulty rather than environmental and social factors. While we detected no significant effects of group size, connectivity, and environmental heterogeneity, larger sample sizes may be needed to reveal more subtle influences on innovation. Together, these results suggest that urbanization gradients can shape behavioral responses to novelty independently from problem-solving abilities.

**Lay Summary:** Cities are challenging places for wildlife, but some species, like sulphur-crested cockatoos, adapt by finding new ways to feed and solve problems. We studied how urbanization and social factors vary with cockatoos’ ability to innovate when presented with extractive-foraging tasks. We found that birds in more urban areas approached the tasks more quickly, suggesting reduced neophobia or increased familiarity with novel objects. This work highlights behavioral differences across urbanization gradients in human-dominated environments.

## Introduction

The question of why some species and individuals thrive in cities while others are negatively affected has become a major focus in behavioral ecology (Lee and Thornton, 2021). Cities represent highly novel habitats, exposing animals to cognitive challenges and opportunities such as new food sources or stimuli, rapid change, and unpredictability (Sayol et al., 2020; Sol et al., 2005). In response, many species have invented a range of solutions to urban opportunities, with examples ranging from using cars to crack nuts in large-billed crows (*Corvus macrorhynchos*) (Nihei and Higuchi, 2002), to tits (*Cyanistes caeruleus* and *Parus major*) piercing milk bottles to forage on cream (Aplin, 2016; Fisher, 1949). Such animal innovations, defined as novel behavioral patterns that address new challenges or present new solutions to familiar problems (Reader and Laland, 2003), have been proposed as being a potentially important component of urban adaptation. In support of this, evidence suggests that urban-dwelling species tend to be more innovative than their rural counterparts (Audet et al., 2016; Møller, 2009; Preiszner et al., 2017), although results remain mixed (Griffin and Guez, 2014; Papp et al., 2015; Vincze and Kovács, 2022).

At the individual level, innovation is a multi-component behavior (Prasher et al., 2019). At the initial point of approach, individual variation in motivation, neophobia and/or neophilia will influence likelihood of interacting with a problem (Greenberg and Mettke-Hofmann, 2001; Mettke-Hofmann, 2007). This will be further influenced by social context, with individuals in groups often facilitated to approach tasks through the benefits of shared vigilance, or experiencing increased motivation with social competition (Miller et al., 2022). Then once an animal engages with the problem, problem-solving success is predicted by a combination of behavioral factors, including previous experience, motor-diversity and persistence (Griffin and Guez, 2014). Finally, cognitive abilities may also influence problem-solving performance, for example via causal reasoning and flexibility (Tebbich et al., 2016).

The social and physical environment can influence each of these factors, leading to population-level variation in problem-solving performance. For instance, larger groups may contain a greater diversity of skills or experiences needed to solve a problem, termed the ‘skill-pool effect’ (Giraldeau, 1984; Morand-Ferron and Quinn, 2011; Prasher et al., 2019). Problem-solving may also be more likely in larger groups due to a non-linear relationship between group size and solving probability (Cantor et al., 2020). Novel and changing habitats may filter for bolder individuals that may be more likely to interact with novel stimuli (Lee and Thornton, 2021; Mettke-Hofmann, 2014), and may also provide individuals with prior experience of interacting with novelty, leading to a reduction in neophobia. Such habitats may also provide a range of diverse resources that promote motor skill learning (Griffin, 2016). Finally, environments characterized by physical or temporal heterogeneity can act as a form of natural enrichment, offering diverse stimuli that influence cognitive development (Braithwaite and Salvanes, 2005; Bredy et al., 2003; Gro Vea Salvanes et al., 2013; Grove, 2012; Rojas-Ferrer and Morand-Ferron, 2020).

In general, and perhaps unsurprisingly given this complexity, studies examining variation in problem-solving abilities between urban and less urban populations have produced mixed results (Audet et al., 2016; Johnson-Ulrich et al., 2021; Møller, 2009; Papp et al., 2015). This may partly reflect the fact that most studies have either examined variation in individuals or populations through a simplified binary comparison or linear gradient from urban to rural/wilderness (Griffin and Guez, 2014; Johnson-Ulrich and Forss, 2025), (but see Chow et al. (2021)). In addition, individuals are often also isolated for testing individually, either in the wild (e.g., Johnson-Ulrich et al. (2021)) or by being brought into the lab (e.g., Morand-Ferron et al. (2011)). While valuable, these approaches have two potential issues. First, urban environments vary in multiple dimensions that affect socio-ecological conditions that, in turn, shape individual innovativeness, with these dimensions often not fully captured by a binary or gradient site comparison (Chow et al., 2021, 2024; Lee and Thornton, 2021). Second, observed innovation rates in populations may be influenced by a combination of individual innovativeness and population demographics (Cantor et al., 2020). Higher-resolution environmental and social data, combined with wild experiments, is thus required to disentangle these effects.

Here we aim to disentangle environmental and social characteristics influencing the likelihood of innovation in wild sulphur-crested cockatoos (*Cacatua galerita*, hereafter SC-cockatoos), by comparing group-level variation in performance on an foraging puzzle across an urban matrix. All roost sites were located within Canberra’s urban-to-suburban matrix, ranging from low-density suburban areas with extensive tree cover to high-density urban centers. SC-cockatoos are common across urban and suburban areas in Australia (Aplin et al., 2021; Davis et al., 2012; Smeele et al., 2022), where several innovative behaviors have been documented (Klump et al., 2021, 2025). Importantly for the purposes of this study, SC-cockatoos have a discrete social structure whereby individuals gather together in communal night roosts, with birds exploiting a shared home range around this roost (Penndorf et al., 2023). This allows for accurate estimates of local group size, as well as high resolution mapping of the environmental conditions experienced by each group, thus allowing us to investigate how variation in urbanization intensity and social density influences innovation.

We installed extractive-foraging tasks that require innovative problem-solving of varying difficulty (Cole et al., 2011; Prasher et al., 2019) in trees at fifteen roost sites around the city of Canberra. We then measured group size (roost size and connectivity) and constructed indices of urbanization and environmental heterogeneity for the area around each roost, comparing these metrics to the time to approach and solve each task. We hypothesized that both environmental and social factors would influence the likelihood of approach and problem-solving across sites. First, when considering the time to approach, we predicted that larger groups and more urban roosting-groups would approach tasks faster, due to a higher probability of chance encounters in larger groups, and due to urban individuals having lower neophobia and/or more familiarity with novel objects (Lee and Thornton, 2021). However, we expected no difference in approach with task difficulty, and had no a-priori predictions for environmental heterogeneity. Second, we predicted that the lower-difficulty tasks would be solved faster. We further predicted faster solving in larger groups due to the non-linear group size effect (Cantor et al., 2020), and faster solving in groups experiencing more environmental heterogeneity due to the effect of environmental complexity on cognitive development (Gro Vea Salvanes et al., 2013; Grove, 2012; Rojas-Ferrer and Morand-Ferron, 2020). Finally, we predicted that once it was disentangled from other co-varying factors, there would still be an direct effect of urbanization on solving, either due to a selective filter on local population composition or due to urban birds’ greater experience interacting with objects requiring diverse motor actions.

## Methods

### Study system

The study was conducted on a population of sulphur crested cockatoos (*Cacatua galerita*) in Canberra, Australia. SC-cockatoos form communal sleeping roosts all year round, aggregating at dusk in groups that range from ten to hundreds of individuals. These roosts split into small foraging sub-groups during the day, generally foraging in a shared home-range over an approximate 2-3 km radius around the roost (Aplin et al., 2021; Fehlmann et al., 2024). Individuals will also roost-switch, sleeping at neighbouring roosts (Penndorf et al., 2023).

The research was carried out across the northern and the western urban and suburban area of Canberra (Figure 1E). Across this area, we identified 27 SC-cockatoo roost sites, and selected 15 focal roost sites for the experiment (see Table 1 in *Supplementary material*).

**Figure 1:**
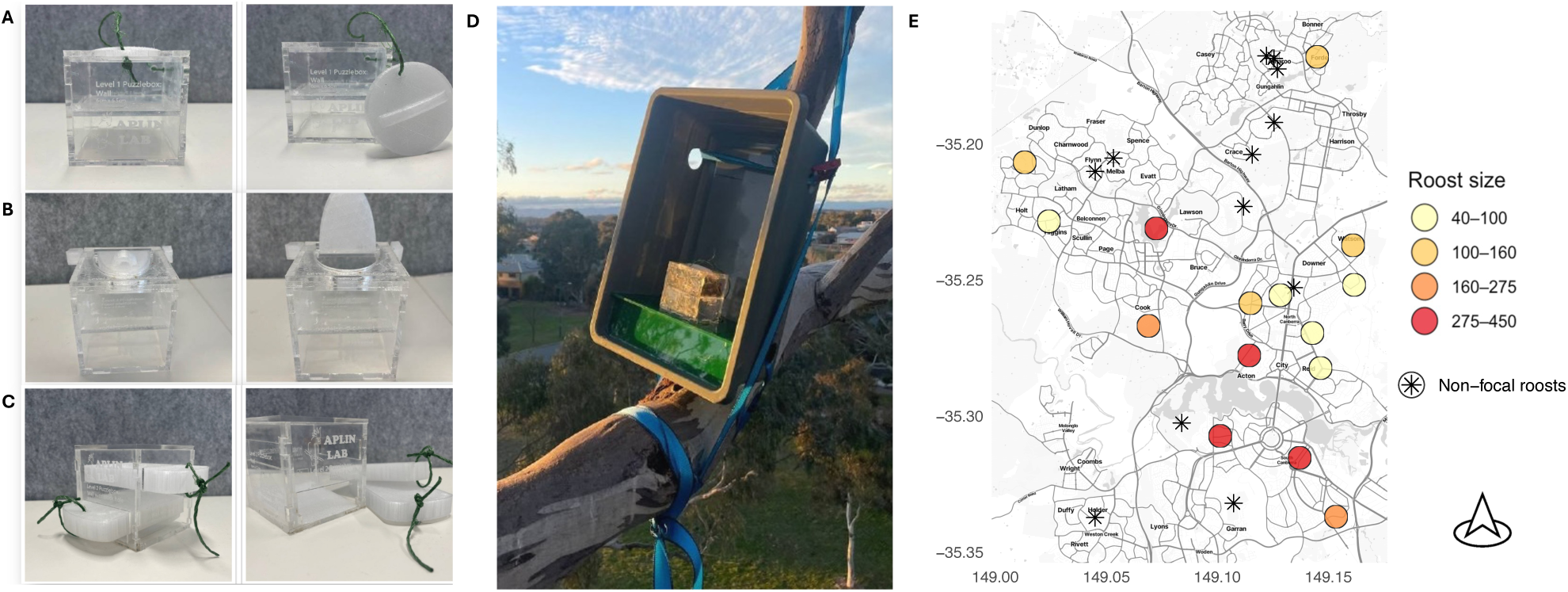
**(A-C)**. Innovative problem-solving tasks shown in unsolved and solved states, presented in order of increasing difficulty from (A) Level 1 - removing lid, (B) Level 2 - moving lever to lift lid, to (C) Level 3-removing two levers such that food drops to opening at bottom. All tasks were baited with approximately 100g of sunflower seed. **(D)**. Level 1 task installed on a tree at Gungahlin, Canberra (ACT), showing support structure and attachment. **(E)**. Known roosts in the urban region of Canberra. Coloured circles indicate the 15 focal roosts included in the experiment; colours represent the estimated roost size, grouped into four categories based on quartile breaks. Black stars represent non-focal roosts, which were not tested but were included in the extended population sizes.

### Experimental design

We tested innovation using food-motivated tasks that required problem solving (Cole et al., 2011; Prasher et al., 2019; Van Horik and Madden, 2016). These comprised 8x8x8 cm custom-made clear acrylic boxes, which were made by laser cutting 3 mm transparent acrylic sheets, and 3D-printed moving components in transparent poly-carbonate. We made three different types of tasks based on low, medium and advanced complexity of motor action (Figure 1). Level 1 consisted of a box containing approximately 100g of sunflower seed, with a removable lid (Figure 1A). Level 2 consisted of the same box as for level 1, but was solved by moving a lever on the side to raise the lid (Figure 1B). Level 3 consisted of two shelves that needed to be removed to access the seed (Figure 1C) (Morand-Ferron and Quinn, 2011). The boxes were glued inside a support structure (Figure 1D) consisting of a grey 38x25x15 cm box. This was then fixed to trees with pull-tie-down straps (Figure 1D).

The experiment was conducted between 14/06 and 24/10/2023 during the Austral winter/early spring. Three tasks, one each per level of difficulty, were installed on trees at each of the 15 roost sites, for a total of 45 tasks across the study population. Roosting sites often consisted of a line of trees (personal observation), and we ascertained all trees used by the roost at a dawn or dusk count prior to installation. The tasks were then positioned randomly within the set of roosting trees to the west, east, or center of the roost in relation to geographic north (see Table 1 *Supplementary material*). All set-ups were located at approximately two-thirds of the tree height, with the distance from the ground varying based on the tree height. Each box was monitored with a HyperFire 2™ camera trap (Reconyx Inc.) positioned on the same branch approximately 1 m away. The camera was triggered by motion detection, and an integrated infrared sensor allowed the camera to record activity even in low-light conditions. After installation at a given roost site, the time was recorded and the experiment began (see Table 2, *Supplementary material*). The puzzle boxes then remained in place for a maximum of 8 weeks, or until they were found solved.

## Social and Environmental metrics

### Local population size

At each roosting location, we quantified local population size, defined as the number of individuals sleeping at a given roost. A roost-count for each site was performed on the evening or morning after puzzle boxes installation by either counting the animals leaving the site at sunrise or entering it at dusk (see Table 2 in *Supplementary material*). In five cases it was not feasible to count roost size on the day of installation due to adverse weather conditions. For those sites, roost size was imputed as the median between counts made closest in time before and after the installation date, using data collected as part of another lab project (see Table 2 in *Supplementary material*). Overall, roost sizes varied from 41 (Lyneham) to 454 (Lake Ginninderra), with a median of 165 birds (see Table 2 in *Supplementary material*).

### Extended population size

As individuals are known both to visit and overlap in foraging home range with nearby roosts, the effective population size might also include the neighbouring roosts. We therefore included a measure of roost connectivity within the wider roosting social landscape as a proxy for extended population size. This was calculated as weighted network centrality (Farine and Whitehead, 2015), where network edges were calculated for each node as the number of other roosts within 2.5 km radius, including all known roosts (focal and non-focal).

### Urbanization index

We described the urban environment surrounding each roost using a composite metric of urbanization derived from land cover and building characteristics. Land cover data were derived from the European Space Agency (ESA) WorldCover 2020 product, which provides data at 10 m resolution based on Sentinel-1 and Sentinel-2 imagery (https://esa-worldcover.org/en). This dataset includes 11 land cover classes: tree cover, shrubland, grassland, cropland, built-up, bare/sparse vegetation, snow and ice, permanent water bodies, herbaceous wetland, mangroves, and moss and lichens. For our analysis, we combined Shrubland, Cropland, and Bare/Sparse vegetation into a single category labelled ‘Vegetation’. We then calculated the proportion of area within a 2 km radius around each roost location that was covered by trees, grass, vegetation, and urban features. We decided to employ a 2 km radius, equivalent to an area of 12.57 km², which approximately matched the average home range size observed during the same field season as part of another project (*unpublished data*).

We described buildings using the 2024 ‘Buildings © Geoscape, Australia’ dataset, which includes digital representations of buildings larger than 9 m² derived from remotely sensed imagery. For each 2 km radius area surrounding a roost, we calculated the average building height and area, as well as the proportion of residential buildings.

To assign an urbanization index to each site, we performed a Principal Component Analysis (PCA), following the method used in Fehlmann et al. (2024). This PCA included a set of variables describing land cover and built features (average building height, average building area and building use) at each site. We used the first principal component from this analysis as our urbanization index, where negative values represent low urbanization and positive values represent high urbanization. This represents the relative variation in anthropogenic modification across sites, capturing the intensity or urban modification from a structural and ecological perspective. As a reference for low urbanization, we included a nearby site with minimal human disturbance, located in the Tidbinbilla Nature Reserve (coordinates: −35.49149, 148.8247).

Vegetation-related variables were included in our PCA to ensure that the index captured habitat transformation that is associated with urban sprawl. Although we refer to our metric as an *urbanization index*, it captures physical and quantifiable features of the environment likely to influence animal behavior and does not include aspects of urbanization such as noise, pollution, population density or socio-economic factors. The index should therefore be viewed as a compound measure that reflects the extent of human development and the associated loss of natural habitat.

The urbanization index varied substantially across the 15 focal roost sites (range: −1.23 to 1.41; mean ± SD = 0.39 ± 0.80), showing considerable variation in urbanization intensity within Canberra’s urban-to-suburban matrix. This range spanned from low-density suburban sites characterized by extensive tree cover and minimal built infrastructure to high-density urban centers dominated by tall buildings and impervious surfaces.

### Environmental heterogeneity

We measured environmental heterogeneity using the O’Neill’s Entropy, which provides the residual amount of entropy in the landscape once the spatial information has been accounted for. We prepared the layer for the analysis in QGIS (QGIS Geographic Information System, Open Source Geospatial Foundation Project, http://qgis.org, version 3.30.2). We rasterized the 2024 *‘Buildings* © *Geoscape, Australia’* layer using the rasterize tool in QGIS, maintaining the original resolution of the shapefile layer. We included roads from the publicly available Microsoft Road Detections (MS roads) dataset, created using automatic image recognition from Bing Maps imagery between 2020 and 2022. We rasterized the layer using the rasterize tool in QGIS, preserving the original resolution. Finally, we merged The European Space Agency (ESA) WorldCover 10 m 2020, the building raster and road raster created as described. We then calculated O’Neill’s entropy for a 2 km radius area around each roost site (O’Neill et al., 1988). We performed the analysis in R Studio (version 4.3.3) using the R-Package *SpatEntropy* (Altieri et al., 2021).

For each site we obtained an absolute measure of entropy and a relative measure of entropy. As these measures were highly correlated (Pearson’s *r* = 0.94, 95% CI [0.83, 0.98], *t*(13) = 9.94, *p* < 0.001), we choose to report and use only the absolute measure of entropy.

### Statistical analysis

To investigate the relationship between time to first approach or solving and our social and environmental variables, we employed Cox proportional-hazards mixed models using the packages *survival*, *survminer*, and *coxme* in Rstudio (version 2024.04.2-764), (Anderson et al., 2021; McCune et al., 2025). We implemented two models: model 1 examined time (in minutes) to first approach, and model 2 examined time to solving. In the main text, we report results based on solving time calculated from the time of first approach, however, we also ran equivalent models considering solving time from initial installation, which produced qualitatively identical results (see *Supplementary material*). We considered urbanization, environmental entropy, roost size, roost centrality, and task difficulty as fixed predictors in both models, and included roost ID as random factor to account for non-independence of observations within roosts. Effects were considered statistically significant when *p* ≤ 0.05.

When no approach or no solving occurred during the experimental period, times were right-censored using the time of puzzle box removal. In the case of missing data due to camera failure, the end point was considered the last available video frame; if no data was recorded at all, the end point was considered 1 minute after the installation time. In the case in which a non-target species solved the puzzle-box, the time for Model 2 was censored at the point of solving by that species.

To complement the time-to-event analysis and assess whether environmental and social factors influenced the probability of solving independent of time, we fitted a third model (Model 3) using binomial logistic regression. This model examined solving success (solved by SCC = 1, not solved by SCC = 0) as a binary outcome, with the same fixed predictors (urbanization, environmental entropy, roost size, roost centrality, and task difficulty) plus time to first approach as an additional covariate.

Given that our field-based approach differs from traditional laboratory problem-solving studies in temporal scale and measurement, we conducted additional sensitivity analyses by including in the models engagement time (cumulative interaction time) as an alternative metric more comparable to controlled experiments. These analyses are detailed in the *Supplementary material*.

To confirm our results, we repeated the analysis in Bayesian framework, using the *brms* package (Brard et al., 2017; Bürkner, 2017; Farhin and Khan, 2022). We included the same fixed effect as before, as well as a random intercept for roost ID. The three models were fitted using four chains of 20000 iterations each (5000 warmup) with an *adapt_delta* of 0.999, and weakly informative priors for all coefficients (normal(0, 1) for fixed effects and the intercept, and an exponential(1) for the standard deviation of the random effect).

### Ethical note

Experimental procedures complied with Australian law and were approved by the Australian National University Animal Experimentation Ethics Committee (Animal Ethics protocol number A2023/05). All fieldwork was conducted under an Australian Capital Territory scientific licence (license number LT20236).

## Results

### Model 1: Time to Approach

When considering the time to first approach of the tasks, the strongest predictor was urbanization, with more urbanized roosts approaching the tasks significantly faster (β = 0.79, hazard ratio [HR] = 2.21, *p* = 0.03,: Figure 2A). The complementary Bayesian analysis supported this findings, with 95% of the posterior distribution lying above 0 (see *Supplementary material*). None of the other predictors had a significant effect: environmental heterogeneity (β = −0.27, hazard ratio [HR] = 0.76, *p* = 0.49: Figure 2C), task difficulty (level 2: β = −0.84, hazard ratio [HR] = 0.43, *p* = 0.09; level 3: β = −0.80, hazard ratio [HR] = 0.45, *p* = 0.15: Figure 2E), roost size (β = −0.06, hazard ratio [HR] = 0.94, *p* = 0.86), or roost connectivity (β = 0.99, hazard ratio [HR] = 2.69, *p* = 0.12) - see supplementary material for the full set of figures.

**Figure 2:**
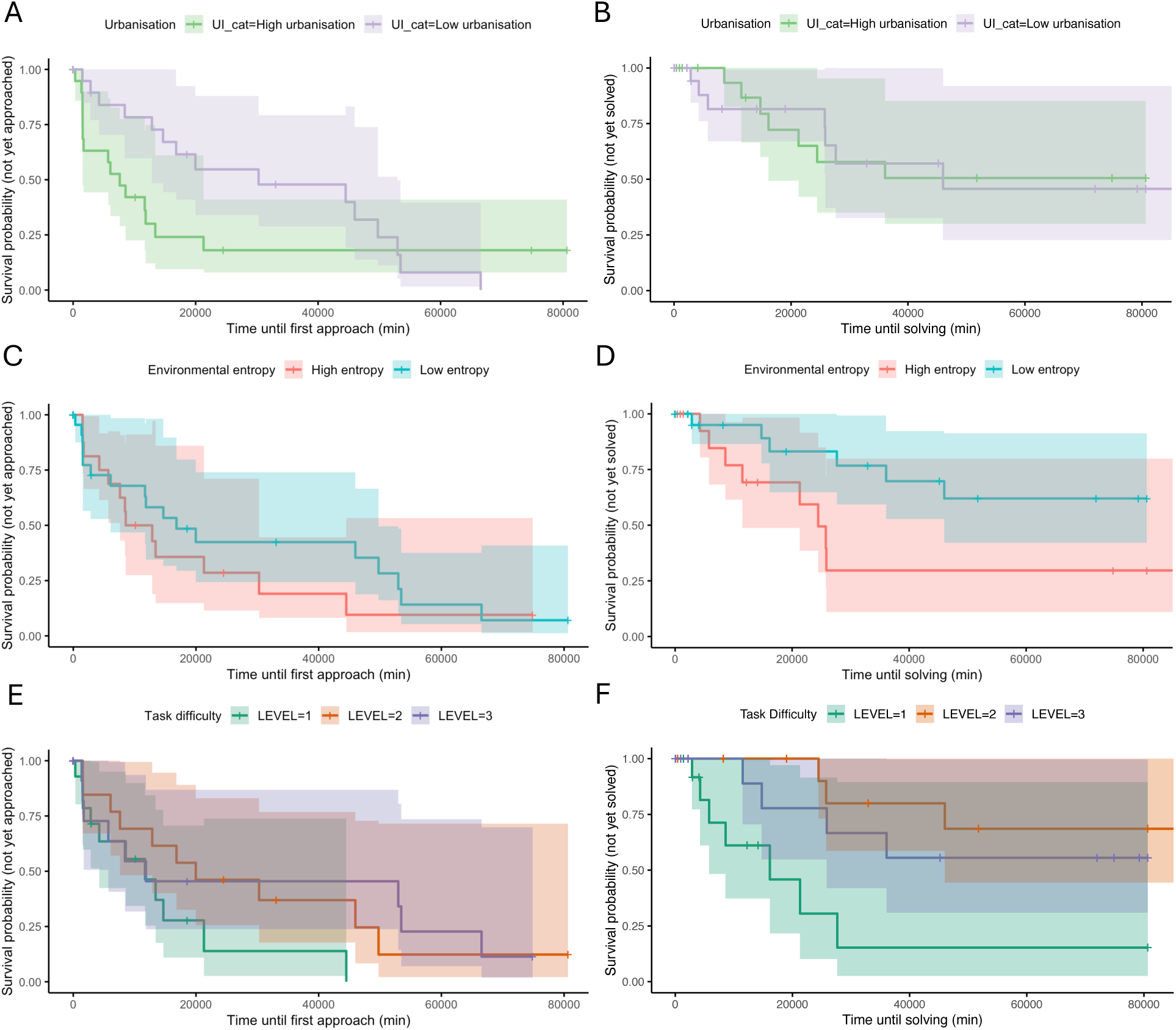
Kaplan–Meier survival curves showing probability of approach (A,C,E) or solve (B,D,F) by a SC-cockatoo across 15 roosts and 45 tasks. **(A–B)**. Probability that SC-cockatoos had not yet approached (A) or solved (B) the task, grouped by urbanization level (green = high [above median], purple = low [at or below median]). SC-cockatoos in more urban areas were significantly faster to approach (*p* = 0.035), but there was no difference in time to solve (*p* = 0.84). **(C-D)**. Probability of not yet approaching (C) or solving (D) the task, grouped by environmental heterogeneity (red = high [above median], azure = low [at or below median]). There was no difference in time to approach (*p* = 0.49), but a non-significant trend for SC-cockatoos in environments of higher heterogeneity to solve tasks faster (*p* = 0.09). **(E-F)**. Probability of not yet approaching (E) or solving (F) the task, grouped by task difficulty (green = Level 1, orange = Level 2, violet = Level 3). Level 1 tasks were solved significantly faster than Level 2 (*p* = 0.0018) or Level 3 (*p* = 0.011).

### Model 2: Time to Solve

When considering the time to first solve, the strongest predictor was task difficulty. As predicted, the level 1 task was solved significantly faster than the other levels (level 2: β = −2.58, hazard ratio [HR] = 0.08, *p* = 0.002; level 3: β = −1.99, hazard ratio [HR] = 0.14, *p* = 0.01: Figure 2F).

Similarly to Model 1, there was no effect of roost size (β = 0.15, hazard ratio [HR] = 1.16, *p* = 0.71) or connectivity (β = 0.85, hazard ratio [HR] = 2.34, *p* = 0.26). However, contrary to Model 1, there was no effect of urbanization (β = −0.09, hazard ratio [HR] = 0.91, *p* = 0.84: Figure 2B). There was a non-significant trend toward faster solving in environments with higher environmental heterogeneity (β = 0.86, hazard ratio [HR] = 2.36, *p* = 0.09, Figure 2D), and the complementary Bayesian analysis indicated that 90% of the posterior distribution supported this effect (see *Supplementary material*).

In both models, Schoenfeld residuals indicated no violation of the proportional hazards assumption overall (first approach model: *p* = 0.12; solving model *p* = 0.24).

### Model 3: Probability of Solving

In contrast to the time-based analyses, none of the predictors significantly influenced solving probability: task difficulty (level 2: β = −1.52, odds ratio [OR] = 0.22, *p* = 0.16; level 3: β = −0.74, OR = 0.48, *p* = 0.48), approach time (β = −0.13, OR = 0.88, *p* = 0.80), roost size (β = −0.31, OR = 0.73, *p* = 0.61), connectivity (β = 0.51, OR = 1.66, *p* = 0.60), urbanization (β = −0.06, OR = 0.94, *p* = 0.92), or environmental heterogeneity (β = 0.74, OR = 2.09, *p* = 0.27). Complementary Bayesian analyses yielded consistent results (see *Supplementary material*).

## Discussion

Urban environments have long been hypothesized to promote innovation in animals (Reader and Laland, 2003; Sol et al., 2013). However empirical studies have produced mixed results, potentially because urban environments can vary at local scales in multiple ways that have both direct and indirect effect on behavior (Chow et al., 2021, 2024; delBarco Trillo and Putman, 2023; Garden et al., 2006; Johnson-Ulrich and Forss, 2025; Lee and Thornton, 2021). To disentangle factors influencing innovativeness in urban-living sulphur-crested cockatoos, we installed problem-solving tasks in roosts at range of urban locations, and measured both time to approach and solve the puzzles. Importantly, this species is a communally roosting central-place forager (Fehlmann et al., 2024; Penndorf et al., 2023), meaning that variation in the experienced social and physical environments can be estimated from discrete locations. We were therefore able to vary our tasks and locations along three axes: task difficulty, local population size, and habitat structure (built environment and heterogeneity). Our results showed that, as predicted, urbanization influenced SC-cockatoos’ initial response to novelty, with individuals in more urban roosts approaching the tasks faster. However, this effect did not extend to task-solving performance, which was influenced by task difficulty with a non-significant trend for an additional effect of environmental heterogeneity. Contrary to expectation, social factors as measured in our study did not predict either approach or solving time. Below, we consider each of these in turn.

### Urbanization

The urbanization metric we have employed in our study primarily reflects the density and the scale of built infrastructure associated with loss of natural habitat. Model 1 results therefore suggest that such areas with higher building density may be occupied by individuals with lower levels of neophobia (fear of novelty (Greenberg and Mettke-Hofmann, 2001)). The relationship between urbanization and neophobia varies across species, with studies reporting reduced neophobia in some urban populations, no differences in others, or even increased neophobia in highly disturbed areas (Griffin et al., 2017). Our results align with studies finding reduced neophobia in urban environments. For example, urban house sparrows (*Passer domesticus*) approach artificial feeders faster than the rural counterpart (Tryjanowski et al., 2016), while a comparative study in crows (*Corvidae*) found that species that exploit the urban environment show less neophobia when exposed to novel objects (Miller et al., 2023).

An increase in building height and density generally also reflects an increase in human population size and human-derived objects. Therefore, another possible explanation for our result may be that individuals may not differ in neophobia *per se*, but rather that individuals living in more urban habitats are more habituated to human-derived objects that resembled our task (Griffin, 2016). As an extension of this, individuals living closer to the city centre may also have had more previous knowledge of the food reward we employed in the experiment (sunflower seeds) (De León et al., 2019; Morisseau and Caro, 2025). These explanations invoke within-individual plasticity through learning or habituation. However, an alternative mechanism is that urban environments selectively favor or attract individuals that are inherently bolder or less neophobic, leading to non-random behavioral sorting across the urbanization gradient (Mazza et al., 2021). We cannot disentangle these explanations in our study, but our results suggest spatial co-variance between urbanization and neophobia is a promising direction for future work in this species.

We further hypothesized that urbanization would have a direct effect on innovative problem-solving, with more urban birds having more access to, and prior experience with, objects involving diverse motor skills. However against our predictions, we didn’t detect any relationship between local variation in urbanization and time to solve. As we didn’t measure motor skills directly, it might be that our premise is incorrect, and there is no correlation between motor skills and urbanization in this species. Alternatively, our tasks may not have required complex motor skills to solve. While we consider this unlikely given the significant effect of task level on solving time, our observations of solving were not detailed enough to allow us to disentangle motor skills from other cognitive influences on success such as persistence or causal reasoning. Additionally, with only 15 roosts, we cannot rule out that we lacked adequate power to detect an effect of urbanization on solving performance, particularly if such an effect is weaker than the strong influence of urbanization on approach behavior.

Overall, our results therefore suggest that urbanization correlates with variation in the behavioral state of SC-cockatoos, with individuals at more urbanized sites showing greater boldness, neophilia, or exploratory behavior, but no difference in other motor or cognitive abilities associated with problem-solving (Morton et al., 2023; Vincze et al., 2024). This contrasts with studies reporting enhanced problem-solving in urban populations (Audet et al., 2016; Møller, 2009; Preiszner et al., 2017), but aligns with findings showing that urbanization can affect motivation or neophobia without necessarily enhancing cognitive performance (Mazza et al., 2021; Morton et al., 2023). For example, in red foxes (*Vulpes vulpes*), urbanization positively affected the approach to, but not the exploitation of, novel food (Morton et al., 2023). SC-cockatoos eat a large variety of food types across their range, many of which involve extractive foraging and fine-scale manipulation of hand, tongue and beak, for example to extract seeds from gum-nuts. It is possible that in common with other highly generalist species, their dietary niche has largely pre-adapted them to the full range of motor skills they might need to exploit urban environments.

While our study did not include truly rural populations, our urbanization index captured substantial variation in environmental conditions within Canberra’s urban matrix (UI range: −1.23 to 1.41, SD = 0.80). This within-city approach complements traditional urban-rural comparisons by examining how fine-scale gradients in urbanization intensity influence behavior (Chow et al., 2021; Fehlmann et al., 2024). This continuous measure of urbanization may be more relevant for understanding behavioral plasticity in urban-adapted species than binary rural-urban classifications, as animals experience and respond to gradual environmental gradients rather than discrete categories. However, our findings should be interpreted with caution when comparing to studies that contrast urban versus rural populations, as we cannot address whether the patterns we observe within cities also apply across the broader urban-rural spectrum.

### Environmental heterogeneity

Intriguingly, unlike for our metric of urbanization, we observed a non-significant trend for environmental heterogeneity on time to solve, with the hazard ratio for solving increasing as heterogeneity decreased. Several factors may have limited our ability to detect a significant effect. First, our sample size (n = 15 roosts) may have provided insufficient statistical power to detect moderate effects. Second, the environmental variation within our study area may not have been sufficient to capture the full range of heterogeneity effects. While urbanization and environmental heterogeneity are often assumed to covary, this relationship is not consistent across contexts (Botequilha Leitao and Ahern, 2002). In our study, urbanization and environmental heterogeneity were not strongly correlated, meaning that the indices measured distinct properties of the environment with heterogeneity reflecting structural complexity independently from the broader urban development.

If the observed marginal trend reflects a real effect, it could align with previous literature positing that environmental variability has been a key driver of encephalization and problem-solving (Grove, 2012; Gro Vea Salvanes et al., 2013; Sol et al., 2005). Further work examining problem-solving across a larger sample size and a broader range of environmental variation is needed to confirm whether the observed trend represents a genuine ecological pattern.

### Social influences

We didn’t observe any effect of local population size or roost connectivity on either time to approach the tasks or to solve them. Our first finding is in contrast to our hypothesis that larger groups would be more likely by chance to encounter the task, and more likely to approach due to the dual effects of lowered predation risk (Shaffer et al., 2021) and increased competition for resources (Griffin and Guez, 2015). One possible explanation for this might be social inhibition, for example as shown in Atlantic salmon (*Salmo salar*) (Brown and Laland, 2002), where individuals may hesitate to engage with tasks due to interference competition (Griffin et al., 2013). Another possibility is social negotiation, for example as shown in ravens (*Corvus corax*) (Stöwe et al., 2006). Here individuals adopt a ‘wait-and-see’ strategy where they monitor the behavior of others before deciding to interact with the task themselves (Griffin et al., 2013). Alternatively, it might be that the effect of group size operates at a level not captured by our measure, for example, overall roost size might not be indicative of the numbers of individuals using the exact tree where each task was installed, or in the group present at the time of encounter. If our measure of local population size did not reflect group sizes at this highly localized level, this may also explain why we didn’t observe any impact of roost size and connectivity on time to solve. Finally, our sample size (n = 15 roosts) may have provided insufficient statistical power to detect moderate effects of social factors. Given the variation in roost sizes and connectivity patterns across our study area, larger sample sizes would be needed to disentangle these complex social dynamics and their potential influence on innovation.

### Conclusion

Urbanization can change conditions in multiple ways that can influence observed innovation rates. Urban-adapted animals often live in higher social densities in urban areas, potentially decreasing neophobia and increasing problem-solving probabilities through social facilitation, group-size effects and increased skill pools. In addition, urban environments contain novel objects and resources that may decrease neophobia, increase neophilia, and improve motor skills (Chow et al., 2025). Finally, some urban environments may exhibit relatively higher environmental heterogeneity at more local scales, with a positive effect on cognitive performance (Tebbich et al., 2016; Griffin and Guez, 2014). Disentangling these influences remains a major challenge for the field (Chow et al., 2021; Reader and Laland, 2003). In this study, we integrated high-resolution continuous measures of urbanization and environmental entropy with discrete measures of local group size and connectivity, allowing us to explore how fine-scale variation in habitat structure and demography might influence innovation rates. Our results suggest that urbanization influenced approach behavior but not solving time in our study population, consistent with some previous findings showing dissociations between motivation and problem-solving performance. However, given our study’s limitations — including sample size, and the challenges of measuring problem-solving in free-ranging populations — further research across broader spatial scales and diverse taxa is needed to determine whether behavioral responses to novelty consistently covary with urbanization independently from cognitive problem-solving abilities. These results underscore the need for ecologically valid studies that incorporate both environmental and social dimensions, and set ground for future research on how cognitive traits are expressed in dynamic, human-modified landscapes.

## Acknowledgements

Thank you to Daniel Noble for the advice on statistical analysis, and to the *Di*fl*cult Birds Research Group* for lending the Reconyx camera traps. Thank you to Fraser Campbell, MacLean Cobden, and Rene Riedelbauch for their help in the data collection, especially during tree climbing. We also thank F. Campbell for help in building the cognitive tasks, and ANU makerspace for providing facilities to build the equipment. We acknowledge the Ngunnawal and Ngambri traditional custodians of the land on which this study was conducted, and pay our respects to their Elders past and present.

## Data availability

All data and code, including STL and Adobe files for 3D-printing and laser cutting respectively, are available on the public GitHub repository https://github.com/LisaFontana96/Innovation.

