## Supplementary for "Social and ecological factors associated with innovation in urban sulphur-crested cockatoos (*Cacatua galerita*)"

### Supplementary Material

#### Study population

##### GPS location and installation date for all roosts

| Roost | GPS | Level 1 GPS | Level 2 GPS | Level 3 GPS | Installation date | Inst. end time |
|---|---|---|---|---|---|---|
| ANU | -35.28, 149.11 | -35.2773, 149.1143 | -35.2786, 149.1139 | -35.2779, 149.1140 | 29/06/2023 | 14:05 |
| CK | -35.27, 149.07 | -35.2665, 149.0696 | -35.2668, 149.0694 | -35.2669, 149.0690 | 24/06/2023 | 10:30 |
| CP | -35.27, 149.14 | -35.2690, 149.1419 | -35.2695, 149.1424 | -35.2691, 149.1429 | 15/07/2023 | 16:50 |
| GPM | -35.21, 149.01 | -35.2080, 149.0144 | -35.2082, 149.0146 | -35.2071, 149.0145 | 16/06/2023 | 15:10 |
| GU | -35.17, 149.14 | -35.1678, 149.1449 | -35.1679, 149.1452 | -35.1680, 149.1457 | 14/06/2023 | 16:00 |
| HA | -35.25, 149.16 | -35.2520, 149.1610 | -35.2518, 149.1615 | -35.2511, 149.1627 | 18/07/2023 | 11:45 |
| HIG | -35.23, 149.02 | -35.2287, 149.0242 | -35.2289, 149.0244 | -35.2240, 149.0242 | 20/06/2023 | 13:20 |
| LG | -35.23, 149.07 | -35.2313, 149.0732 | -35.2302, 149.0735 | -35.2317, 149.0728 | 20/06/2023 | 10:40 |
| LY | -35.26, 149.13 | -35.2556, 149.1284 | -35.2539, 149.1275 | -35.2546, 149.1277 | 24/06/2023 | 14:00 |
| NAR | -35.34, 149.15 | -35.3368, 149.1524 | -35.3369, 149.1515 | -35.3374, 149.1539 | 29/08/2023 | 17:00 |
| OC | -35.26, 149.11 | -35.2587, 149.1146 | -35.2589, 149.1139 | -35.2587, 149.1151 | 24/06/2023 | 15:30 |
| TP | -35.31, 149.14 | -35.3141, 149.1380 | -35.3163, 149.1369 | -35.3145, 149.1374 | 16/08/2023 | 11:50 |
| WA | -35.24, 149.16 | -35.2373, 149.1611 | -35.2376, 149.1616 | -35.2371, 149.1609 | 03/07/2023 | 13:15 |
| WM | -35.28, 149.15 | -35.2827, 149.1464 | -35.2838, 149.1452 | -35.2836, 149.1457 | 18/07/2023 | 13:50 |
| YA | -35.31, 149.10 | -35.3069, 149.1010 | -35.3068, 149.1004 | -35.3074, 149.1014 | 24/08/2023 | 11:50 |

Table 1: GPS coordinates and installation date & time for all the levels in all the study sites. ANU- Australian National University, CK- Cook, CP- Corroboree Park, GPM- Garis Place McGregor, GU- Gungahlin, HA- Hackett, HIG- Higgins, LG- Lake Ginninderra, LY- Lyneham, NAR- Narrabundah, OC- O'Connor, TP- Telopea Park, WA- Watson, WM- War Memorial, YA- Yarralulma

### Social variables

#### Roost counts

| Roost site | Count 1 | Date | Count 2 | Date | Count 3 | Date | Count 4 | Date | Median |
|---|---|---|---|---|---|---|---|---|---|
| ANU | 287 | 29/06/23 | NA | NA | NA | NA | NA | NA | NA |
| CK | 176 | 24/06/23 | NA | NA | NA | NA | NA | NA | NA |
| CP | 52 | 15/07/23 | NA | NA | NA | NA | NA | NA | NA |
| GP | 165 | 16/06/23 | NA | NA | NA | NA | NA | NA | NA |
| GU | 130 | 14/06/23 | NA | NA | NA | NA | NA | NA | NA |
| HA | 231 | 18/07/23 | 80 | 13/07/23 | 93 | 06/07/23 | 32 | 20/07/23 | 86.5 |
| HIG | 103 | 20/06/23 | NA | NA | NA | NA | NA | NA | NA |
| LG | 454 | 20/06/23 | NA | NA | NA | NA | NA | NA | NA |
| LY | 25 | 24/06/23 | 56 | 14/07/23 | 41 | 30/06/23 | NA | NA | 41 |
| NAR | 270 | 29/08/23 | NA | NA | NA | NA | NA | NA | NA |
| OC | 136 | 24/06/23 | 193 | 12/07/23 | NA | NA | NA | NA | 164.5 |
| TP | 327 | 16/08/23 | NA | NA | NA | NA | NA | NA | NA |
| WA | 156 | 03/07/23 | 145 | 22/07/23 | NA | NA | NA | NA | 150.5 |
| WM | 28 | 18/07/23 | 138 | 29/07/23 | 107 | 09/07/23 | 57 | 20/07/23 | 82 |
| YA | 280 | 24/08/23 | NA | NA | NA | NA | NA | NA | NA |

Table 2: Count of individuals for each one of the 15 roosts included in the study; numbers in regular text represent the counts performed the same day of the puzzle boxes installation, while the numbers in bold were obtained by computing the mean between counts of different days. ANU- Australian National University, CK- Cook, CP- Corroboree Park, GPM- Garis Place McGregor, GU- Gungahlin, HA- Hackett, HIG- Higgins, LG- Lake Ginninderra, LY- Lyneham, NAR- Narrabundah, OC- O'Connor, TP- Telopea Park, WA- Watson, WM- War Memorial, YA- Yarralulma. Roost counts for the locations HA, LY, OC, WM, and WA were obtained by computing the median among counts done in different days as part of another lab project.

### Environmental heterogeneity and urbanization index - European Space Agency (ESA) WorldCover 2020 product at 10 m resolution

#### Urbanization index

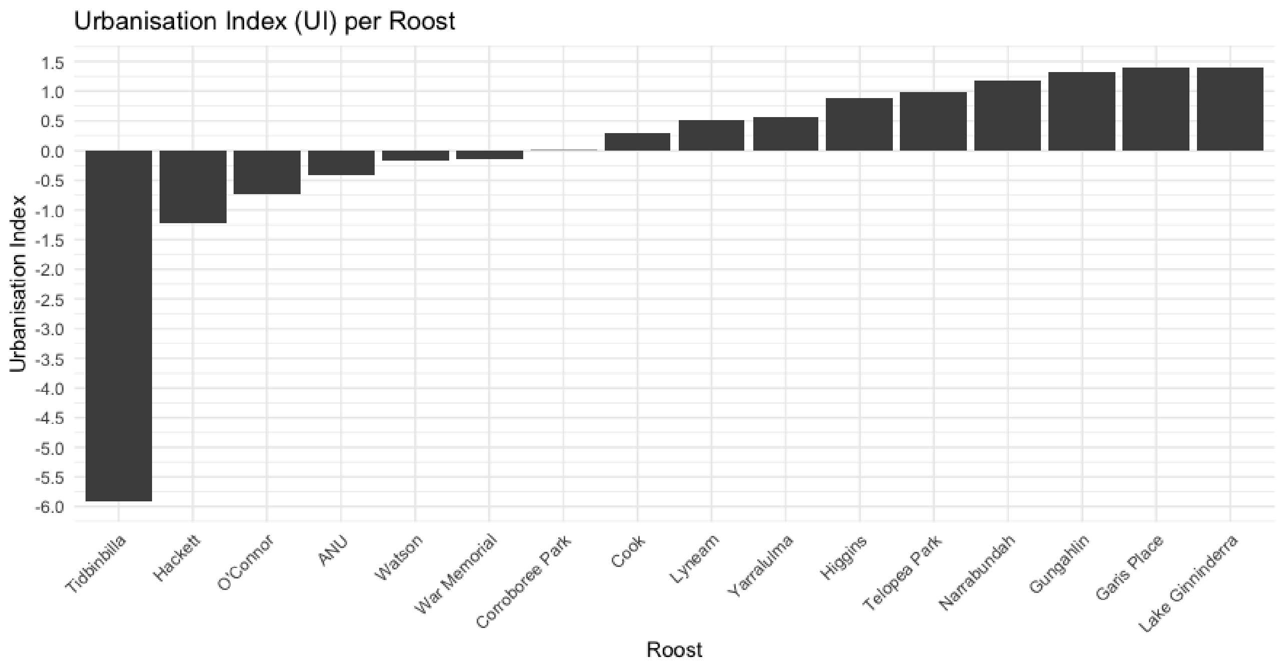


Figure 1: Urbanization index using ESA landcover for the 15 roost sites included in the study and the standard for low urbanization. To provide a standard for low urbanization, we included in the analysis a roost site in an area without human disturbance (Tidbinbilla Nature Reserve; -35.49149, 148.8247).

#### Environmental heterogeneity

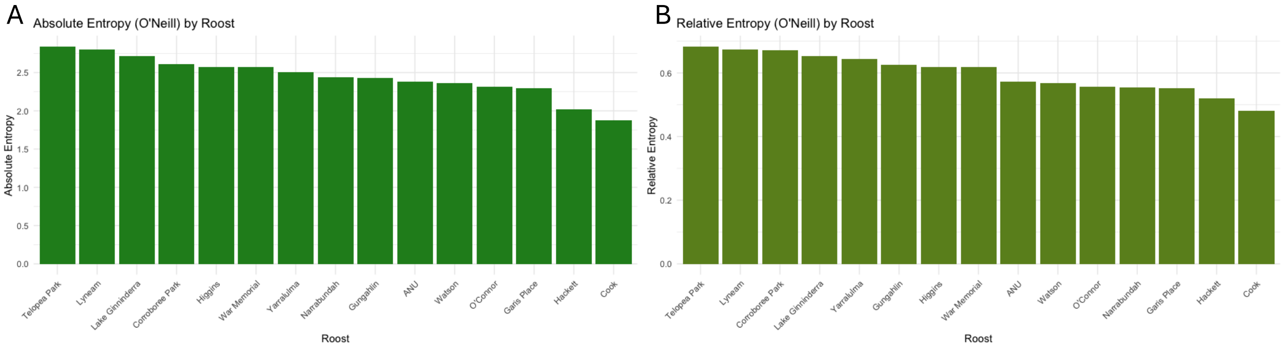


Figure 2: O'Neill index using ESA landcover for the 15 roost sites included in the study
(**A**) Absolute value of O'Neill index (**B**) Relative value of O'Neill index

#### Summary indexes for all roosts

O'Neill and urbanization indexes were weakly positively correlated (Pearson's $r$ = 0.43, 95% CI [0.15, 0.64], $t(43) = 3.01$, $p < 0.003$).

| Roost site | Urbanization index | Absolute O'Neill index | Relative O'Neill index |
|---|---:|---:|---:|
| ANU | -0.411539234 | 2.38556 | 0.573606 |
| CK | 0.307843649 | 1.872539 | 0.4811473 |
| CP | -0.008539533 | 2.612594 | 0.6713038 |
| GPM | 1.397632510 | 2.296369 | 0.5521599 |
| GU | 1.325772421 | 2.431183 | 0.6246905 |
| HA | -1.229035332 | 2.022477 | 0.5196738 |
| HIG | 0.895896786 | 2.56934 | 0.6177957 |
| LG | 1.406180015 | 2.711737 | 0.6520349 |
| LY | 0.506605631 | 2.804285 | 0.674288 |
| NAR | 1.185692518 | 2.439549 | 0.5551433 |
| OC | -0.731595913 | 2.310014 | 0.5554409 |
| TP | 0.991942454 | 2.84118 | 0.6831593 |
| WM | -0.133344554 | 2.569044 | 0.6177246 |
| WA | -0.172817052 | 2.364287 | 0.5684909 |
| YA | 0.568560545 | 2.501617 | 0.6427883 |
| TB | -5.899254911 | NA | NA |

Table 3: Urbanization and entropy indexes for each of the 15 roost sites and the control site (TB = Tidbinbilla).

### Video analysis

Videos collected from the camera traps were played with IINA media-player software for macOS (v. 1.3.5). In addition to SC-cockatoos, other animals occasionally approached the boxes, although SC-cockatoo were the species that provided most approaches. These approaches were documented, but only data from the focal species were considered in the analysis. For SC-cockatoos, each approach, defined as the presence on camera in close proximity to the setup, was recorded along with the corresponding date and time. The start time for an approach was marked by the subject's appearance on camera, while the approach was considered to have ended when the subject was no longer visible for at least 15 seconds. Approaches were categorized as *solving*, *presence* or *attempt*: *presence* indicated proximity to the set-up without interaction, *attempt* indicated clear attention to or active attempt to solve the box, and *solving* indicated actual solving of the puzzle. For approaches categorized as *solving*, the time of access to the reward was also recorded.

#### Task-solving time across roosts by difficulty level

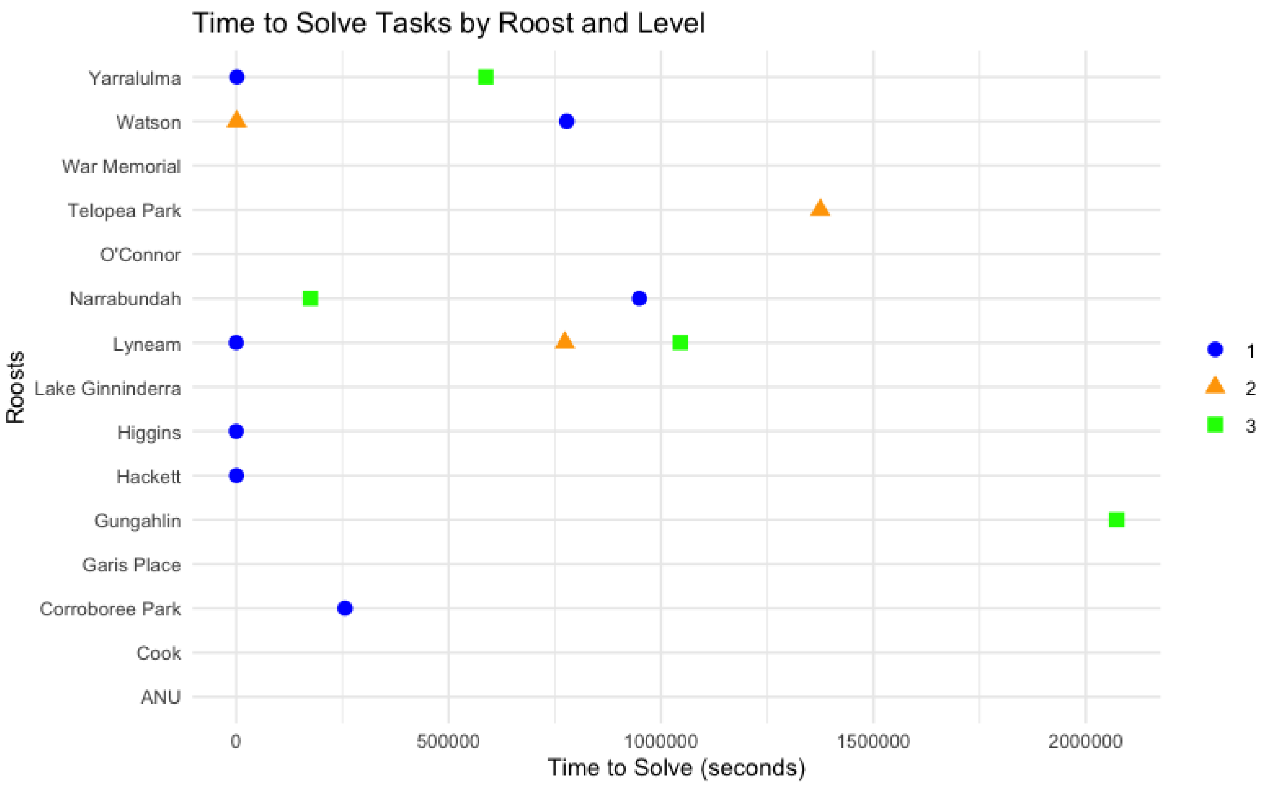


Figure 3: Time to solve each task across roost sites. Each point represents the time taken (in seconds) to solve a task at a given roost. Task difficulty is indicated by shape and colour: Level 1 (easy) — blue circles, Level 2 (intermediate) — orange triangles, Level 3 (hard) — green squares.

#### Success rate per predictor

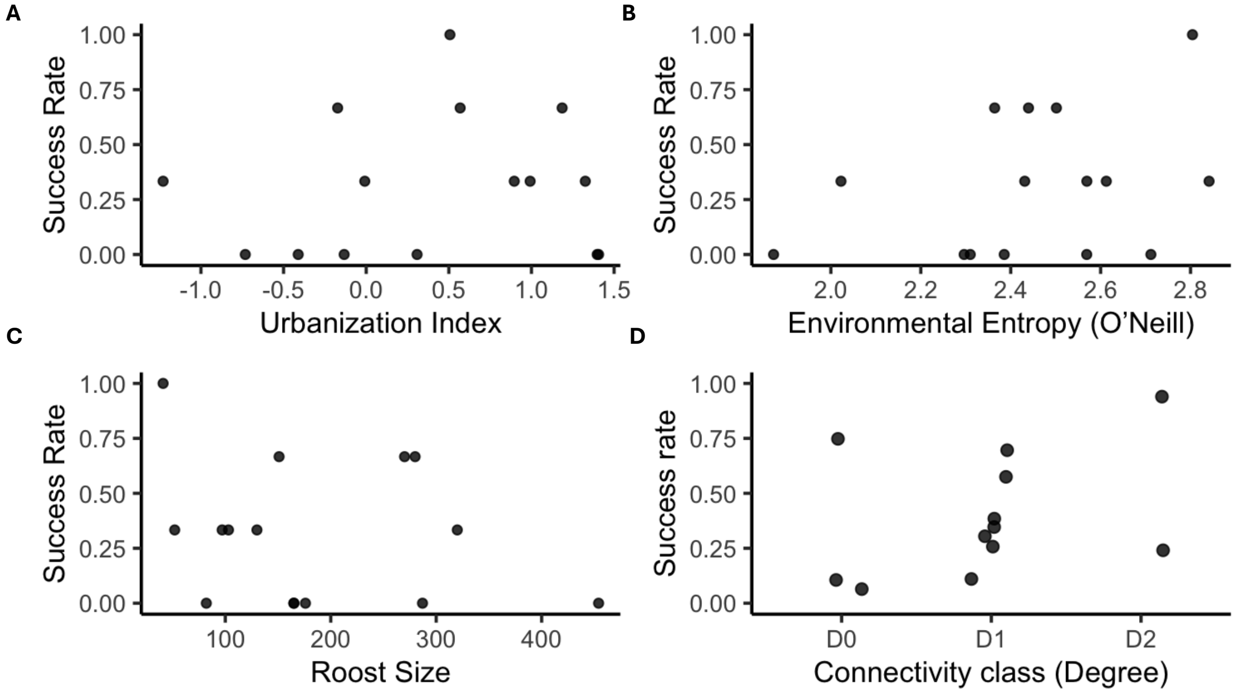


Figure 4: Observed success rate of sulphur-crested cockatoos across roosts plotted against (A) Urbanization Index, (B) Environmental Entropy (O’Neill index), (C) Roost Size, and (D) Connectivity (Degree). Each point represents one roost (n = 15). The figure displays raw, descriptive data. D0 = degree equals 0l; D1 = degree equals 1; D2 = degree equals 2.

### Summary models as presented in the main text

#### Cox proportional hazards model: time to first approach

| Variable | coef | exp(coef) | SE | z | p-value |
|---|---|---|---|---|---|
| LEVEL 2 | -0.840 | 0.432 | 0.494 | -1.70 | 0.089 |
| LEVEL 3 | -0.805 | 0.447 | 0.561 | -1.44 | 0.151 |
| Roost size (scaled) | -0.058 | 0.943 | 0.327 | -0.18 | 0.858 |
| Degree | 0.990 | 2.691 | 0.637 | 1.55 | 0.120 |
| **Urbanization (scaled)** | **0.792** | **2.207** | **0.376** | **2.11** | **0.035** |
| O’Neill entropy (scaled) | -0.273 | 0.761 | 0.395 | -0.69 | 0.489 |
| **Random effects:** | | | | | |
| Roost (Intercept): SD = 0.532 Variance = 0.283 | | | | | |
| **Model fit:** | | | | | |
| Events: 30 N: 45 | | | | | |
| Penalized log-likelihood: 20.58 AIC: 2.52 *p* = 0.0149 | | | | | |

Table 4: Summary of Cox proportional hazards model for time to first approach. Positive coefficients indicate increased hazard (shorter times).

### Cox proportional hazards model: time to solve (from first approach)

| Variable | coef | se(coef) | Chisq | p-value | exp(coef) | 95% CI |
|---|---|---|---|---|---|---|
| **LEVEL 2** | **-2.583** | **0.827** | **9.75** | **0.0018** | **0.076** | **[0.015, 0.382]** |
| **LEVEL 3** | **-1.987** | **0.785** | **6.40** | **0.0110** | **0.137** | **[0.029, 0.639]** |
| Roost size (scaled) | 0.146 | 0.387 | 0.14 | 0.7100 | 1.157 | [0.542, 2.468] |
| Degree | 0.850 | 0.757 | 1.26 | 0.2600 | 2.339 | [0.530, 10.320] |
| Urbanization (scaled) | -0.093 | 0.468 | 0.04 | 0.8400 | 0.911 | [0.364, 2.280] |
| O'Neill entropy (scaled) | 0.858 | 0.512 | 2.81 | 0.0940 | 2.358 | [0.865, 6.433] |
| **Random effects:** | | | | | | |
| Frailty (Roost): Chisq = 0.00, $p$ = 0.96 | | | | | | |
| **Model fit:** | | | | | | |
| Concordance = 0.795 (SE = 0.057) | | | | | | |
| Likelihood ratio test = 17.85 on 6 df, $p$ = 0.007 | | | | | | |

Table 5: Summary of Cox proportional hazards model for solving time. Positive coefficients indicate increased hazard (shorter times).

### Cox proportional hazards model: time to solve (from installation end time)

| Variable | coef | se(coef) | Chisq | p-value | exp(coef) | 95% CI |
|---|---|---|---|---|---|---|
| **LEVEL 2** | **-1.902** | **0.757** | **6.32** | **0.012** | **0.149** | **[0.034, 0.658]** |
| **LEVEL 3** | **-1.475** | **0.711** | **4.30** | **0.038** | **0.229** | **[0.057, 0.923]** |
| Roost size (scaled) | -0.219 | 0.363 | 0.36 | 0.550 | 0.804 | [0.395, 1.637] |
| Degree | 0.459 | 0.669 | 0.47 | 0.490 | 1.583 | [0.427, 5.871] |
| Urbanization (scaled) | 0.013 | 0.405 | 0.00 | 0.970 | 1.013 | [0.458, 2.240] |
| O'Neill entropy (scaled) | 0.635 | 0.442 | 2.06 | 0.150 | 1.887 | [0.793, 4.491] |
| **Random effects:** | | | | | | |
| Frailty (Roost): Chisq = 0.00, $p$ = 0.95 | | | | | | |
| **Model fit:** | | | | | | |
| Concordance = 0.748 (SE = 0.077) | | | | | | |
| Likelihood ratio test = 13.47 on 6 df, $p$ = 0.04 | | | | | | |

Table 6: Summary of Cox proportional hazards model for time to solve (alternative solving time model). Positive coefficients indicate increased hazard (shorter latency).

### Logistic regression: solving probability

We initially included a random effect for roosts, but this was removed due to singular fit (variance = $1.09 \times 10^{-8}$), resulting in a standard logistic regression rather than GLMM.

| Variable | Estimate | SE | z value | p-value | Odds Ratio | 95% CI |
|---|---|---|---|---|---|---|
| Time to 1st approach (scaled) | -0.128 | 0.501 | -0.255 | 0.798 | 0.88 | [0.30, 2.36] |
| Level 2 vs 1 | -1.520 | 1.091 | -1.393 | 0.164 | 0.22 | [0.02, 1.68] |
| Level 3 vs 1 | -0.741 | 1.043 | -0.710 | 0.477 | 0.48 | [0.05, 3.62] |
| Roost size (scaled) | -0.313 | 0.605 | -0.517 | 0.605 | 0.73 | [0.21, 2.41] |
| Degree | 0.509 | 0.974 | 0.523 | 0.601 | 1.66 | [0.24, 12.47] |
| Urbanization (scaled) | -0.061 | 0.581 | -0.105 | 0.917 | 0.94 | [0.29, 3.01] |
| O'Neill entropy (scaled) | 0.739 | 0.669 | 1.105 | 0.269 | 2.09 | [0.59, 8.65] |
| **Model fit:** | | | | | | |
| AIC = 50.6, Null deviance = 41.5, Residual deviance = 34.6 | | | | | | |
| N = 30 (approached tasks only; 15 never-approached tasks excluded) | | | | | | |
| Note: No random effects (standard logistic regression) | | | | | | |

Table 7: Summary of logistic regression for solving probability including time to first approach as predictor. Positive coefficients indicate increased log-odds (higher probability of solving). Note: random effect for roosts was removed due to singular fit (variance ≈ 0).

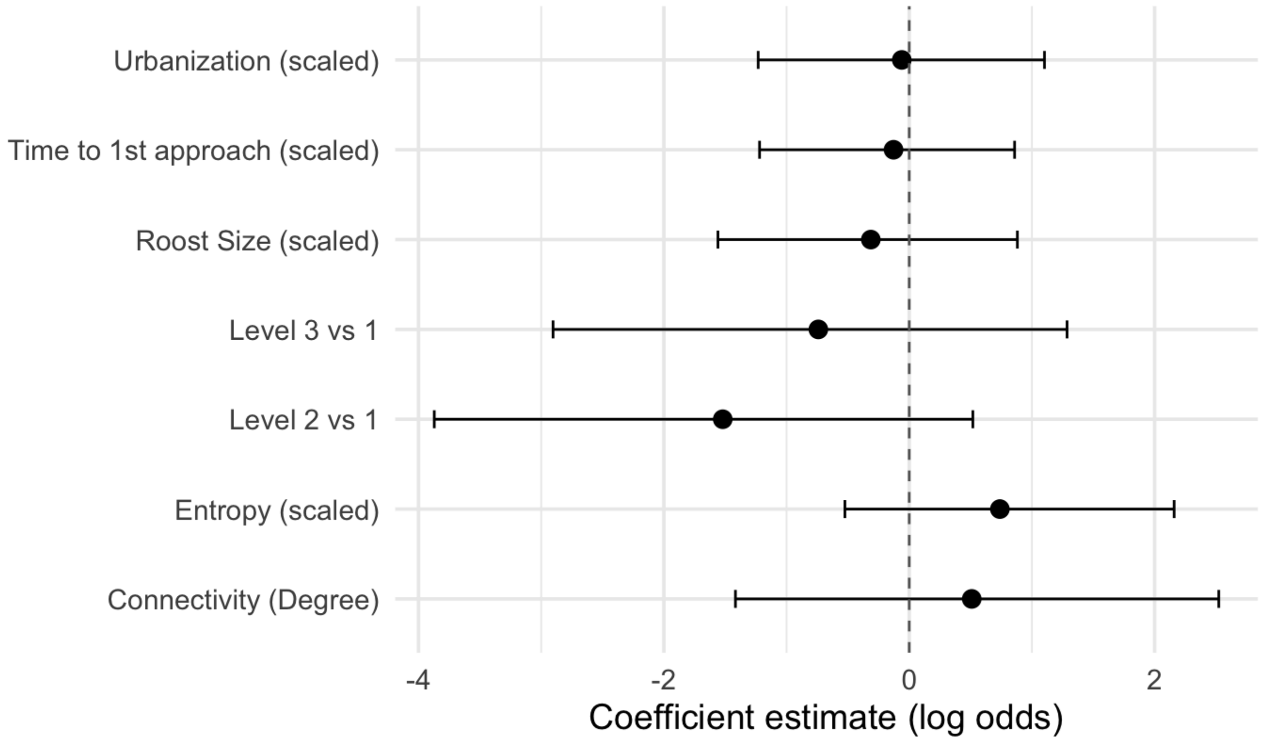


Figure 5: Logistic regression for solving probability including time to first approach as predictor, showing coefficient estimates (log odds) with 95% confidence intervals for each predictor variable.

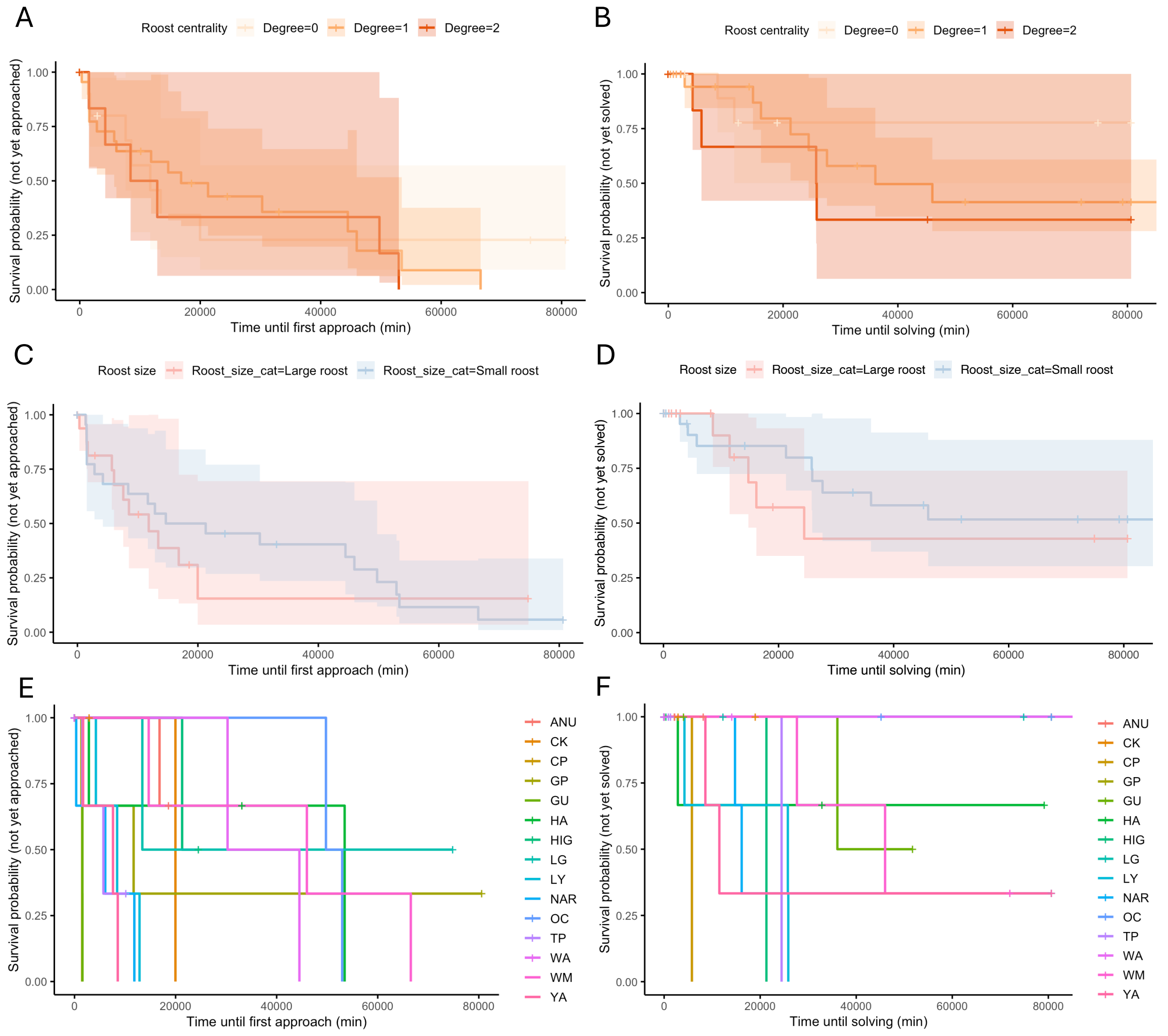


Figure 6: **Kaplan–Meier survival curves showing probability of approach (A,C,E) or solve (B,D,F) by a SC-cockatoo across 15 roosts and 45 tasks. (A–B)** Probability that SC-cockatoos had not yet approached (A) or solved (B) the task, grouped by roost centrality (yellow = degree 0, light orange = degree 1, orange = degree 2). There was no difference in latency to approach ($p = 0.12$) or solving ($p = 0.26$). **(C-D)**: Probability of not yet approaching (C) or solving (D) the task, grouped by roost size (red = large [above median], azure = small [at or below median]). There was no difference in latency to approach ($p = 0.86$) or solving ($p = 0.71$). **(E-F)**: Probability of not yet approaching (E) or solving (F) the task, grouped by roost site. Each line represents a different roost (see legend). Roost identity was included as a random effect in the models, but did not significantly influence approach (variance = 0.238, SD = 0.532) or solving latency ($p = 0.96$).

### Supplementary frequentist models

#### Binomial GLMM: probability of solving; number of approaches

##### Model formula:

```
SOLVED 1/0 ~ LEVEL + scale(Roost size) + Degree + scale(Number of
approaches) + scale(Urbanization) + scale(absolute O'Neill entropy) +
(1 | Roost)
```

| Variable | Estimate | SE | z value | p-value | Odds Ratio | 95% CI |
|---|---|---|---|---|---|---|
| Approaches to solve (scaled) | 0.231 | 0.429 | 0.538 | 0.591 | 1.26 | [0.54, 2.92] |
| Level 2 vs 1 | -1.705 | 1.041 | -1.639 | 0.101 | 0.18 | [0.02, 1.40] |
| Level 3 vs 1 | -1.199 | 0.948 | -1.266 | 0.206 | 0.30 | [0.05, 1.93] |
| Roost size (scaled) | -0.432 | 0.677 | -0.639 | 0.523 | 0.65 | [0.17, 2.45] |
| Degree | 0.397 | 1.152 | 0.344 | 0.731 | 1.49 | [0.16, 14.23] |
| Urbanization (scaled) | 0.191 | 0.676 | 0.283 | 0.777 | 1.21 | [0.32, 4.55] |
| O'Neill entropy (scaled) | 0.629 | 0.757 | 0.831 | 0.406 | 1.88 | [0.43, 8.27] |
| **Random effects:** | | | | | | |
| Roost (Intercept): Variance = 0.906, SD = 0.952 | | | | | | |
| **Model fit:** | | | | | | |
| AIC = 65.5, BIC = 81.8 | | | | | | |
| Log-likelihood = -23.7, Deviance = 47.5 | | | | | | |

Table 8: Summary of binomial GLMM model for solving probability.

### Linear mixed model: predictors of total engagement time

**Model formula:**

```
log(total engagement time) ~ LEVEL + scale(Roost size) + Degree +
scale(Urbanization) + scale(absolute O'Neill entropy) + (1 | Roost)
```

| Variable | Estimate | SE | t value |
|---|---|---|---|
| Level 2 vs 1 | 0.447 | 0.794 | 0.563 |
| Level 3 vs 1 | 0.241 | 0.767 | 0.315 |
| Roost size (scaled) | -0.526 | 0.485 | -1.085 |
| Degree | -1.128 | 0.842 | -1.340 |
| Urbanization (scaled) | -0.095 | 0.480 | -0.198 |
| O'Neill entropy (scaled) | 0.788 | 0.494 | 1.596 |
| **Random effects:** | | | |
| Roost (Intercept): Variance = 0.308, SD = 0.555 | | | |
| Residual: Variance = 2.251, SD = 1.500 | | | |
| **Model fit:** | | | |
| REML criterion = 80.0 | | | |
| N = 24 (approached tasks), Groups: 12 roosts | | | |

Table 9: Summary of linear mixed model examining predictors of total engagement time (log-transformed) for tasks that were approached by cockatoos. Response variable is log(total engagement time in minutes).

### Linear mixed model: predictors of engagement time before solving

**Model formula:**

```
log(total engagement time before solving) ~ LEVEL + scale(Roost size)
+ Degree + scale(Urbanization) + scale(absolute O'Neill entropy) + (1 |
Roost)
```

| Variable | Estimate | SE | t value |
|---|---|---|---|
| Level 2 vs 1 | 0.521 | 0.813 | 0.641 |
| Level 3 vs 1 | 0.293 | 0.782 | 0.374 |
| Roost size (scaled) | -0.504 | 0.511 | -0.988 |
| Degree | -1.123 | 0.890 | -1.262 |
| Urbanization (scaled) | -0.115 | 0.510 | -0.225 |
| O'Neill entropy (scaled) | 0.777 | 0.523 | 1.486 |
| **Random effects:** | | | |
| Roost (Intercept): Variance = 0.435, SD = 0.659 | | | |
| Residual: Variance = 2.329, SD = 1.526 | | | |
| **Model fit:** | | | |
| REML criterion = 81.0 | | | |
| N = 24 (approached tasks), Groups: 12 roosts | | | |

Table 10: Summary of linear mixed model examining predictors of engagement time before solving (log-transformed) for tasks that were approached.

### Binomial GLMM: probability of solving; with engagement time

**Model formula:**

```
SOLVED 1/0 ~ LEVEL + scale(Roost size) + Degree + scale(Urbanization)
+ scale(absolute O'Neill entropy) + scale(total engagement time) + (1 |
Roost)
```

| Variable | Estimate | SE | z value | p-value | Odds Ratio | 95% CI |
|---|---|---|---|---|---|---|
| Engagement time (scaled) | 0.858 | 0.501 | 1.714 | 0.087 | 2.36 | [0.88, 6.29] |
| Level 2 vs 1 | -2.147 | 1.173 | -1.831 | 0.067 | 0.12 | [0.01, 1.16] |
| Level 3 vs 1 | -1.497 | 1.022 | -1.465 | 0.143 | 0.22 | [0.03, 1.66] |
| Roost size (scaled) | -0.153 | 0.723 | -0.212 | 0.832 | 0.86 | [0.21, 3.54] |
| Degree | 0.700 | 1.239 | 0.565 | 0.572 | 2.01 | [0.18, 22.84] |
| Urbanization (scaled) | 0.141 | 0.712 | 0.198 | 0.843 | 1.15 | [0.29, 4.65] |
| O'Neill entropy (scaled) | 0.488 | 0.810 | 0.603 | 0.547 | 1.63 | [0.33, 7.97] |
| **Random effects:** | | | | | | |
| Roost (Intercept): Variance = 1.062, SD = 1.031 | | | | | | |
| **Model fit:** | | | | | | |
| AIC = 61.9, BIC = 78.2 | | | | | | |
| Log-likelihood = -22.0, Deviance = 43.9 | | | | | | |

Table 11: Summary of binomial GLMM model for solving probability with engagement time as predictor.

### Bayesian Cox models

For all the models we used weakly informative priors: normal priors for the fixed effects, normal prior for the intercept, and exponential for the standard error. *Interpretation criteria:* An effect was considered supported when the posterior probability of the coefficient being greater than zero ($P(\beta > 0)$) or less than zero ($P(\beta < 0)$) was greater than or equal to 0.95 (Lin and Yin, 2015; Makowski et al.,

2019). This implies that less than 2.5% of the posterior probability mass lies on the other side of zero, indicating strong evidence for a directional effect, as previously done in other behavioural and ecological research (Ellison, 2004).

#### Bayesian binomial model: probability of solving

**Model formula:**

```
SOLVED 1/0 ~ scale(Approach latency) + LEVEL + scale(Roost size) +
Degree + scale(Urbanization) + scale(absolute O'Neill entropy) + (1 |
Roost)
```

| **Parameter** | **Estimate** | **SE** | **2.5% CI** | **97.5% CI** | **Rhat** | **Bulk ESS** | **Tail ESS** |
|---|---|---|---|---|---|---|---|
| Intercept | -0.51 | 1.24 | -3.18 | 1.84 | 1.00 | 3057 | 2327 |
| Approach latency (scaled) | -0.57 | 0.67 | -2.07 | 0.61 | 1.00 | 2541 | 2163 |
| Level 2 (vs 1) | -1.02 | 0.98 | -3.01 | 0.89 | 1.00 | 4287 | 3188 |
| Level 3 (vs 1) | -0.33 | 0.94 | -2.14 | 1.53 | 1.00 | 4683 | 3257 |
| Roost size (scaled) | -0.70 | 0.79 | -2.38 | 0.74 | 1.00 | 3030 | 2406 |
| Degree | 0.58 | 1.00 | -1.40 | 2.61 | 1.00 | 3447 | 2704 |
| Urbanization (scaled) | -0.04 | 0.75 | -1.47 | 1.48 | 1.00 | 2895 | 2540 |
| O'Neill entropy (scaled) | 0.87 | 0.79 | -0.57 | 2.51 | 1.00 | 3094 | 2891 |
| **Random effect: Roost** | | | | | | | |
| SD(Intercept) = 1.51 (95% CI: 0.07 – 3.91) | | | | | | | |
| N = 30 (approached tasks only; 15 never-approached tasks excluded) | | | | | | | |

Table 12: Summary of Bayesian binomial GLMM for probability of solving including approach latency. Positive coefficients indicate increased log-odds (higher probability of solving).

| **Parameter** | $P(\beta > 0)$ | $P(\beta < 0)$ |
|---|---|---|
| Time to 1st approach (scaled) | 0.192 | 0.808 |
| Level 2 (vs 1) | 0.146 | 0.855 |
| Level 3 (vs 1) | 0.356 | 0.643 |
| Roost size (scaled) | 0.182 | 0.819 |
| Connectivity (Degree) | 0.725 | 0.275 |
| Urbanization (scaled) | 0.475 | 0.525 |
| O'Neill entropy (scaled) | 0.875 | 0.125 |

Table 13: Posterior probabilities ($P(\beta > 0)$ and $P(\beta < 0)$) for fixed effects in the Bayesian binomial GLMM predicting probability of solving including time to first approach.

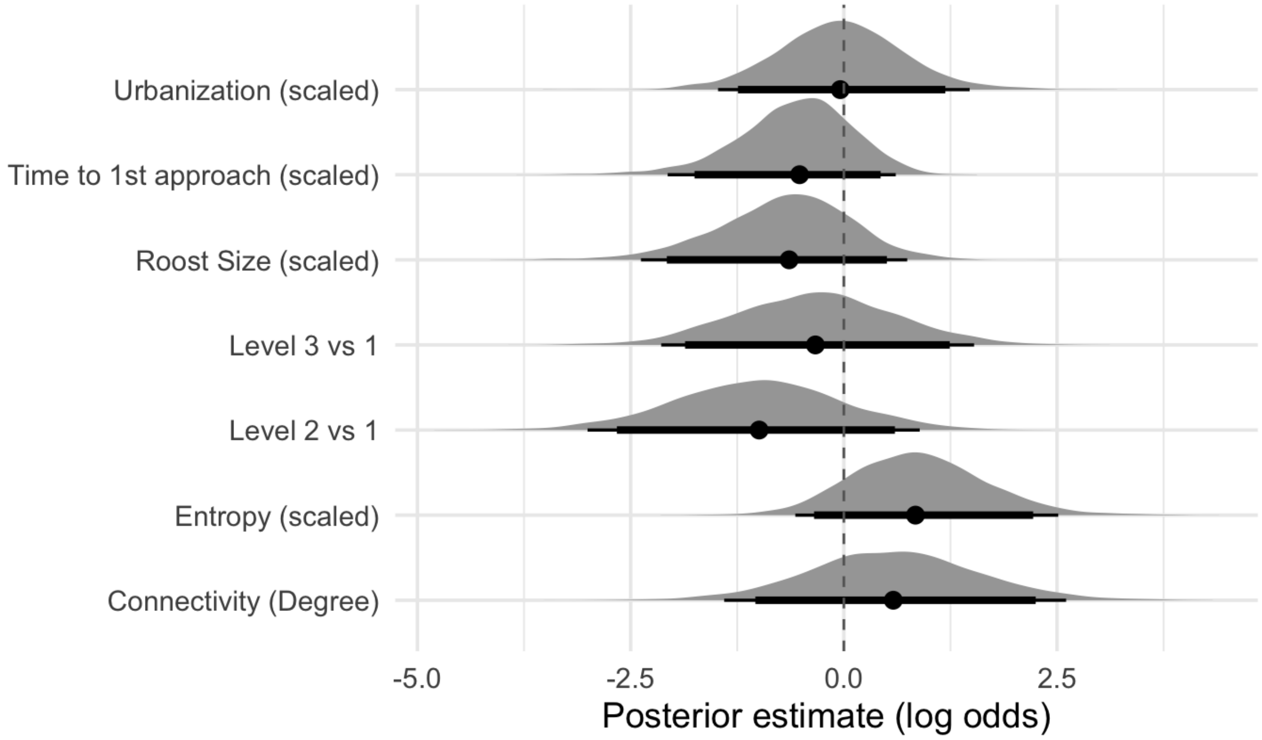


Figure 7: Results of the Bayesian Binomial GLMM (Probability of Solving). Posterior distributions of the regression coefficients (log odds) with 95% credible intervals for each predictor variable.

### Bayesian Cox proportional hazards model: time to first approach

**Model formula:**

```
Surv(time to first approach, censor) ~ LEVEL + scale(Roost size) +
Degree + scale(Urbanization) + scale(absolute O’Neill entropy) + (1 |
Roost)
```

| Parameter | Estimate | SE | 2.5% CI | 97.5% CI | Rhat | Bulk ESS | Tail ESS |
|---|---|---|---|---|---|---|---|
| Intercept | 0.93 | 0.66 | -0.35 | 2.27 | 1.00 | 30742 | 33650 |
| Level 2 (vs 1) | -0.59 | 0.44 | -1.45 | 0.26 | 1.00 | 56871 | 45614 |
| Level 3 (vs 1) | -0.47 | 0.47 | -1.40 | 0.45 | 1.00 | 51089 | 46724 |
| Roost size (scaled) | -0.19 | 0.35 | -0.92 | 0.48 | 1.00 | 32960 | 30311 |
| Degree | 0.55 | 0.56 | -0.59 | 1.62 | 1.00 | 29581 | 34973 |
| Urbanization (scaled) | 0.59 | 0.37 | -0.16 | 1.32 | 1.00 | 34017 | 38073 |
| O’Neill entropy (scaled) | -0.07 | 0.39 | -0.81 | 0.73 | 1.00 | 30868 | 33964 |
| **Random effect: Roost** | | | | | | | |
| SD(Intercept) = 0.71 (95% CI: 0.04 – 1.66) | | | | | | | |

Table 14: Summary of Bayesian Cox model for time to first approach. Positive coefficients indicate increased hazard (shorter latency).

| Parameter | $P(\beta > 0)$ | $P(\beta < 0)$ |
|---|---|---|
| Urbanization (scaled) | **0.954** | **0.046** |
| Connectivity (Degree) | 0.841 | 0.159 |
| O'Neill entropy (scaled) | 0.414 | 0.586 |
| Roost size (scaled) | 0.290 | 0.710 |
| Level 3 (vs 1) | 0.162 | 0.838 |
| Level 2 (vs 1) | 0.089 | 0.911 |

Table 15: Posterior probabilities ($P(\beta > 0)$ and $P(\beta < 0)$) for fixed effects in the Bayesian Cox model predicting time to first approach. Bold values indicate strong directional support (≥ 0.95)

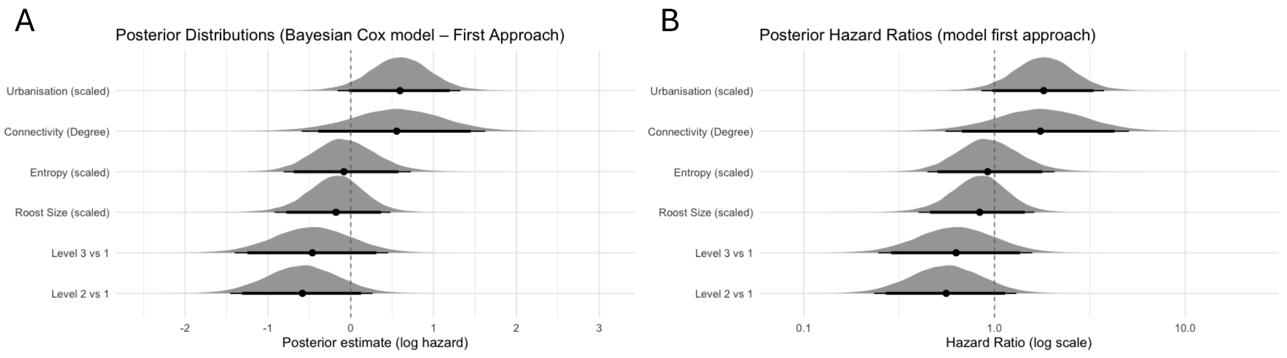


Figure 8: Results of the Bayesian Cox Proportional Hazards Model (First Approach). **A**: Posterior distributions of the regression coefficients (log hazard ratios) for each predictor variable. **B**: Posterior hazard ratios (on a log scale) with 95% credible intervals for each predictor variable.

### Bayesian Cox proportional hazards model: time to solve (from first approach)

#### Model formula:

```
Surv(time to solve, censor) ~ LEVEL + scale(Roost size) + Degree +
scale(Urbanization) + scale(absolute O'Neill entropy) + (1 | Roost)
```

| Parameter | Estimate | SE | 2.5% CI | 97.5% CI | Rhat | Bulk ESS | Tail ESS |
|---|---|---|---|---|---|---|---|
| Intercept | 0.14 | 0.83 | -1.52 | 1.78 | 1.00 | 43441 | 40373 |
| Level 2 (vs 1) | -1.36 | 0.60 | -2.56 | -0.21 | 1.00 | 54650 | 45236 |
| Level 3 (vs 1) | -0.92 | 0.58 | -2.08 | 0.20 | 1.00 | 60134 | 45845 |
| Roost size (scaled) | -0.39 | 0.47 | -1.38 | 0.49 | 1.00 | 38312 | 36509 |
| Connectivity (Degree) | 0.32 | 0.66 | -1.00 | 1.61 | 1.00 | 41667 | 41580 |
| Urbanization (scaled) | 0.09 | 0.48 | -0.83 | 1.06 | 1.00 | 40934 | 37624 |
| O'Neill entropy (scaled) | 0.63 | 0.50 | -0.36 | 1.63 | 1.00 | 39536 | 39759 |
| **Random effect: Roost** | | | | | | | |
| SD(Intercept) = 0.96 (95% CI: 0.04 – 2.43) | | | | | | | |

Table 16: Summary of Bayesian Cox model for time to solve. Positive coefficients indicate increased hazard (shorter times).

| Parameter | $P(\beta > 0)$ | $P(\beta < 0)$ |
|---|---|---|
| Urbanization (scaled) | 0.575 | 0.425 |
| Connectivity (Degree) | 0.694 | 0.306 |
| O'Neill entropy (scaled) | 0.902 | 0.098 |
| Roost size (scaled) | 0.195 | 0.805 |
| Level 3 (vs 1) | **0.045** | **0.955** |
| Level 2 (vs 1) | **0.010** | **0.990** |

Table 17: Posterior probabilities ($P(\beta > 0)$ and $P(\beta < 0)$) for fixed effects in the Bayesian Cox model predicting time to solve. Bold values indicate strong directional support (≥ 0.95)

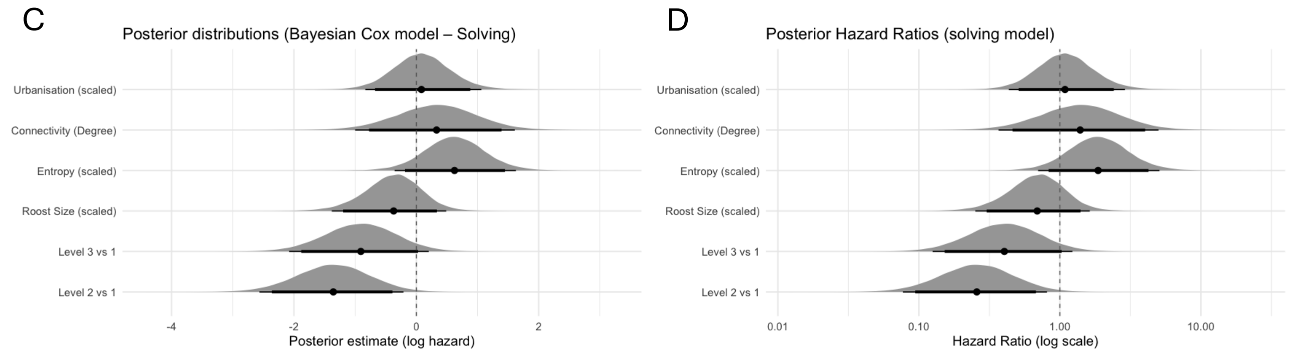


Figure 9: Results of the Bayesian Cox Proportional Hazards Model (Solving). **A**: Posterior distributions of the regression coefficients (log hazard ratios) for each predictor variable. **B**: Posterior hazard ratios (on a log scale) with 95% credible intervals for each predictor variable.

### Bayesian Cox proportional hazards model: time to solve (from installation end time)

**Model formula:**

```
Surv(time to solve 2, censor) ~ LEVEL + scale(Roost size) + Degree +
scale(Urbanization) + scale(absolute O'Neill entropy) + (1 | Roost)
```

| Parameter | Estimate | SE | 2.5% CI | 97.5% CI | Rhat | Bulk ESS | Tail ESS |
|---|---|---|---|---|---|---|---|
| Intercept | 0.10 | 0.76 | -1.43 | 1.55 | 1.00 | 54031 | 44826 |
| Level 2 (vs 1) | -1.43 | 0.58 | -2.61 | -0.30 | 1.00 | 70452 | 46547 |
| Level 3 (vs 1) | -0.85 | 0.57 | -1.96 | 0.26 | 1.00 | 70642 | 47876 |
| Roost size (scaled) | -0.14 | 0.42 | -1.00 | 0.65 | 1.00 | 51492 | 39812 |
| Connectivity (Degree) | 0.50 | 0.61 | -0.71 | 1.69 | 1.00 | 49397 | 46319 |
| Urbanization (scaled) | -0.01 | 0.43 | -0.88 | 0.84 | 1.00 | 50592 | 45387 |
| O'Neill entropy (scaled) | 0.64 | 0.46 | -0.28 | 1.56 | 1.00 | 49040 | 45224 |
| **Random effect: Roost** | | | | | | | |
| SD(Intercept) = 0.64 (95% CI: 0.02 – 1.79) | | | | | | | |

Table 18: Summary of Bayesian Cox model for time to solve (alternative specification). Positive coefficients indicate increased hazard (shorter times).

| **Parameter** | $P(\beta > 0)$ | $P(\beta < 0)$ |
|---|---|---|
| Urbanization (scaled) | 0.488 | 0.512 |
| Connectivity (Degree) | 0.794 | 0.206 |
| O'Neill entropy (scaled) | 0.917 | 0.083 |
| Roost size (scaled) | 0.370 | 0.630 |
| Level 3 (vs 1) | 0.068 | 0.932 |
| Level 2 (vs 1) | **0.007** | **0.993** |

Table 19: Posterior probabilities ($P(\beta > 0)$ and $P(\beta < 0)$) for fixed effects in the Bayesian Cox model predicting time to solve. Bold values indicate strong directional support ($\geq 0.95$).
